# Adverse drug reactions associated with the use of biological agents

**DOI:** 10.1101/2020.09.24.311241

**Authors:** Jorge Enrique Machado-Alba, Anyi Liliana Jiménez-Morales, Yulieth Carolina Moran-Yela, Ilsa Yadira Parrado-Fajardo, Luis Fernando Valladales-Restrepo

## Abstract

**Introduction:** Biotech drugs open new possibilities to treat diseases for which drug therapy is limited, but they may be associated with serious adverse drug reactions (ADRs).

**Objective:** To identify the ADRs associated with the use of biotech drugs in Colombia.

**Methods:** This was a retrospective study of ADR reports from 2014 to 2019, contained in the database of the pharmacovigilance program of Audifarma SA. The ADRs, groups of associated drugs, and affected organs were classified.

**Results:** A total of 5,415 reports of ADRs associated with biotech drugs were identified in 78 Colombian cities. A total of 76.1% of the cases corresponded to women. The majority were classified as type A (55.0%) and B (28.9%), and 16.7% were serious cases. The respiratory tract was the most commonly affected organ system (16.8%), followed by the skin and adnexa (15.6%). Antineoplastic and immunomodulatory drugs accounted for 70.6% of the reports, and the drugs related to the greatest number of ADRs were adalimumab (12.2%) and etanercept (11.6%).

**Conclusions:** There has been an incremental increase in the reporting of ADRs associated with the use of biotech drugs in the pharmacovigilance program, related to the strengthening and appropriation of the patient safety culture and improvement in the quality of the generated information. It is important to empower physicians and entire health teams to ensure the traceability of ADRs and to perform interdisciplinary interventions derived from pharmacovigilance at the individual and population levels.

## Introduction

Biotech drugs are synthesized from expressed proteins, monoclonal antibodies, vectors (viruses, lipid molecules), antibody fragments, antisense molecules and lipid vectors using innovative genetic engineering methods and recombinant DNA technology, which converts them into drug complexes during manufacturing and these drugs offer the potential to participate in important biological processes in humans [1]. Side effects and adverse drug reactions (ADRs) are events that can seriously affect the health of individuals who take drugs for therapeutic, diagnostic or prophylactic purposes. Very often, hospital care may be required due to the presentation of undesirable effects, which may also be responsible for significant mortality [2].

The development and use of biotech drugs is booming in most countries, since these drugs open new possibilities for the treatment of diseases for which drug therapy is limited [3, 4]. They constitute a therapeutic innovation, which has also represented an unknown world of adverse reactions and events that affect patient safety. For this reason, it is necessary to analyze patient records in order to identify all undesirable events and detect early signs that reduce patient risk and to make comparisons with safety profile reports available in international reference entities so that public warnings can be issued [5]. In addition to endangering the health of individuals, ADRs cause treatment abandonment and unexpected costs that affect the finances of health systems, so their early identification can help prevent and solve these problems [6, 7]. Hence, pharmacovigilance is the cornerstone in monitoring drug safety during clinical use [8].

Because information on the safety associated with the use of biotech drugs, the incidence rates of events and their severity, the causality association and the data on the true benefit/risk ratio are often insufficient, our objective was to identify the ADRs related to the use of biotech drugs in patients affiliated with the Colombian Health System between 2014 and 2019.

## Materials and methods

A retrospective study was conducted to analyze the systematized databases of reports of ADRs and suspected ADRs occurring between January 1, 2014, and December 31, 2019, that were associated with the biotech drugs dispensed by the company Audifarma SA. Audifarma is a drug-dispensing logistics operator that covers more than 8.5 million users of the Colombian Health System, corresponding to 17.3% of the population affiliated with it, including patients under the contributory or employer-paid regime and the state-funded regime.

The reports are usually made by the treating physicians, nurses responsible for patient care, administrative personnel involved in treatment adherence monitoring or patient support programs and pharmaceutical chemists in charge of pharmacotherapeutic monitoring of ADR reports. The information was processed by the group of pharmaceutical chemists from Audifarma who receive the reports of suspected ADRs, check the data, input them into the system and analyze each report. In addition, support is provided by a pharmacoepidemiologist when needed. Because the data are typed into the database by different professionals at the national level, the recorded data were checked and verified, and specific compilations were created for annual periods from 2014 to 2019. All of the cases received are included in the pharmacovigilance program and reported to the National Pharmacovigilance Program of the National Institute of Drug and Food Surveillance of Colombia (Instituto Nacional de Vigilancia de Medicamentos y Alimentos - INVIMA) within the established deadlines, including the information required by current legislation.

Only the records of patients with complete information, case follow-up and causality analysis were included. Incomplete records or records considered null were excluded. The grounds for annulment included the following: 1. Report without associated ADR. 2. Duplicate report. 3. Medication not dispensed by Audifarma. 4. Corresponds to a medication error. 5. Corresponds to a quality or nonconforming product complaint. 6. Lack of dates.

The general database included the filing date of the ADR report, city, drug (generic name), drug anatomical therapeutic chemical (ATC) classification (letter code and two digits), severity (serious, not serious), type of ADR according to the Rawlins and Thompson classification (A, B, C, D, E, F), ADR probability classification according to the WHO (definitive, probable, possible, unlikely, conditional, unassessable), reported event and the traceability of report submission to INVIMA.

The reported ADRs were standardized according to the WHO adverse reaction terminology (WHO-ART). The main drugs for which ADRs were reported were classified, describing the first 15 ATC subgroups (letter code and first two digits), and an ADR list was created for the 10 drugs with the highest numbers of reports.

The statistical package SPSS 26.0 for Windows (IBM, USA) was used for data analysis, and the data are expressed as frequencies, percentages and means. Incidence rates were estimated from the ADR reports and total patients who were dispensed biotech drugs per monitoring year and per 100,000 health system affiliates. The present study was approved by the Bioethics Committee of the Universidad Tecnologica de Pereira under the risk-free research category (approval number 0104-2019). The principles established by the Declaration of Helsinki were respected. No personal data of the patients were used.

## Results

A total of 5,415 ADR reports associated with the use of 71 biotech drugs were identified throughout the six years of monitoring, in 78 Colombian cities, and with respect to 10 health insurance companies and 65 healthcare institutions, including clinics and hospitals; a progressive increase in the number of cases was observed (Table 1). A total of 4,122 (76.1%) reports corresponded to female patients.

**Table 1.** Number of reports per year, classification according to severity and type of adverse reactions in patients treated with biological agents in Colombia 2014-2019

|  | Number of cases (n=5500) | Percentage |
| --- | --- | --- |
| **Year of report** |  |  |
| 2014 | 187 | 3.5 |
| 2015 | 211 | 3.9 |
| 2016 | 360 | 6.6 |
| 2017 | 864 | 16.0 |
| 2018 | 1321 | 24.4 |
| 2019 | 2472 | 45.7 |
| **Classification according to severity** |  |  |
| Not serious | 4192 | 77.4 |
| Serious | 903 | 16.7 |
| Lethal | 6 | 0.1 |
| Therapeutic failure | 317 | 5.9 |
| Not classified | 9 | 0.2 |
| **Type of adverse reaction** |  |  |
| A (Augmented) | 2925 | 55.0 |
| B (Bizarre) | 1563 | 28.9 |
| C (Chronic) | 152 | 2.8 |
| D (Delayed) | 87 | 1.6 |
| E (End of treatment) | 4 | 0.1 |
| F (Foreign) | 209 | 3.9 |
| Without classification | 1145 | 21.1 |

According to the ADR severity, 77.4% were classified in the nonserious category, followed by serious events, and six were associated with a fatal outcome. In addition, a low percentage (0.2) could not be classified (Table 1). The drugs associated with lethal ADRs were abatacept (four cases), etanercept (one case) and rituximab (one case). The most common ADR type was type A, followed by type B and type C reactions (Table 1).

According to the ATC classification, antineoplastics and immunomodulators were the groups with the highest number of reports, followed by medications for the respiratory and skeletal muscle systems (Table 2). The therapeutic subgroups most frequently associated with ADRs were immunosuppressants, other antineoplastic agents (including monoclonal antibodies) and drugs for systemic use for obstructive airway diseases (Table 2).

**Table 2.** Classification of biological agents associated with adverse drug reactions according to anatomical therapeutic chemical (ATC) group and subgroup. Colombia 2014-2019.

| ATC | Description according ATC group | Patients | Percentage |
| --- | --- | --- | --- |
| **L** | Antineoplastic and immunomodulating agents | 3823 | 70.6 |
| **R** | Respiratory system | 662 | 12.2 |
| **M** | Musculo-skeletal system | 351 | 6.5 |
| **A** | Alimentary tract and metabolism | 265 | 4.9 |
| **S** | Sensory organs | 137 | 2.5 |
| **J** | Antiinfectives for systemic use | 88 | 1.6 |
| **H** | Systemic hormonal preparations, excl. Sex hormones and insulins | 52 | 1.0 |
| **B** | Blood and blood forming organs | 27 | 0.5 |
| **V** | Various | 4 | 0.1 |
| **C** | Cardiovascular system | 3 | 0.1 |
| **D** | Dermatologicals | 3 | 0.1 |
| **Description according ATC subgroup** |  |  |  |
| **L04A** | Immunosuppressants | 3201 | 59.1 |
| **L01X** | Other antineoplastic agents | 562 | 10.4 |
| **R03D** | Other systemic drugs for obstructive airway diseases | 637 | 11.8 |
| **M05B** | Drugs affecting bone structure and mineralization | 351 | 6.5 |
| **H05A** | Parathyroid hormones and analogues | 1 | 0 |
| **A16A** | Other alimentary tract and metabolism products | 204 | 3.8 |
| **A10A** | Insulins and analogues | 61 | 1.1 |
| **J06B** | Immunoglobulins | 88 | 1.6 |
| **L03A** | Immunostimulants | 60 | 1.1 |
| **S01L** | Ocular vascular disorder agents | 136 | 2.5 |
| **H01A** | Anterior pituitary lobe hormones and analogues | 51 | 0.9 |
| **B02B** | Vitamin k and other hemostatics | 24 | 0.4 |
| **R05C** | Expectorants, excl. Combinations with cough suppressants | 25 | 0.5 |
| **S01E** | Antiglaucoma preparations and miotics | 1 | 0 |
| **C10A** | Lipid modifying agents | 3 | 0.1 |

ATC: anatomical therapeutic chemical

The most common ADRs were those causing respiratory system disorders, followed by skin and adnexal disorders and general disorders (Table 3). The causality analysis determined that most ADRs were considered possibly associated with the reported drug (67.9% were definitive, probable or possible) (Figure 1). The biotech drugs with the highest number of reports in the monitoring period were adalimumab, etanercept and omalizumab. The drugs whose incidence increased the most between 2014 with 2019 were denosumab (30.0% increase), omalizumab (18.4%) and etanercept (15.2%). There was an increased incidence of ADR reports for secukinumab between 2016 and 2019, estimated at 41.7%. There were smaller increases for abatacept (3.2%) and rituximab (1.9%). The total number of reports for each of the 10 biotech drugs with the highest numbers of ADRs, the percentages they represented of all notifications, the incidences per 100 patients who received them and the incidences compared by 100,000 affiliates between 2014 and 2019 are shown in Table 4.

**Table 3.**
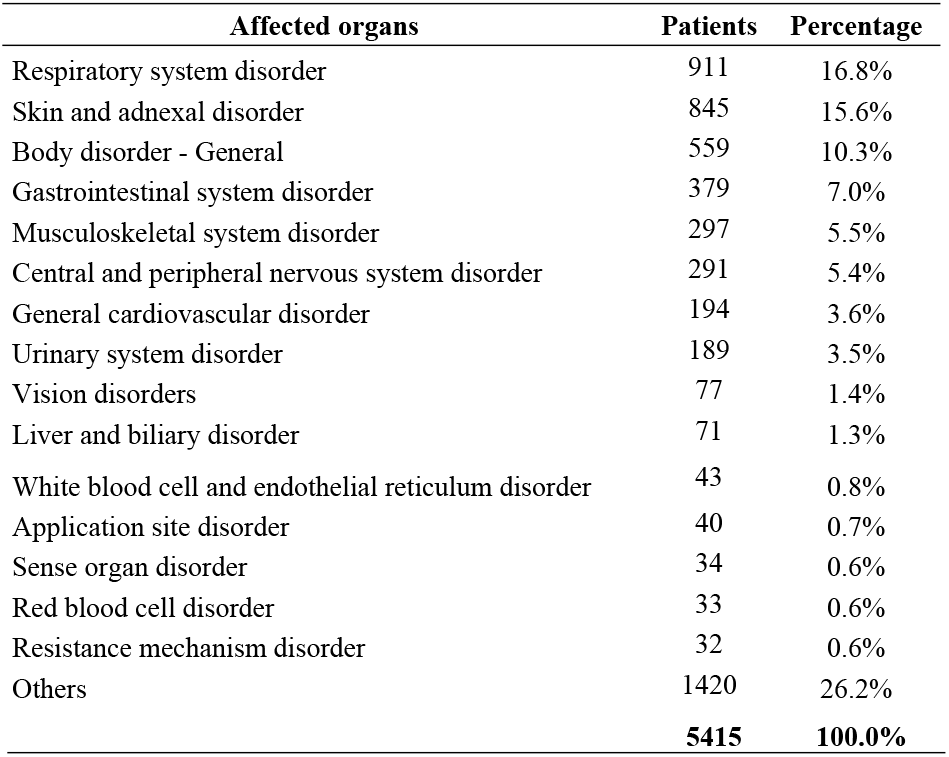
Main systems affected with adverse drug reactions for biological agents in Colombia 2014-2019..

**Figure 1.**
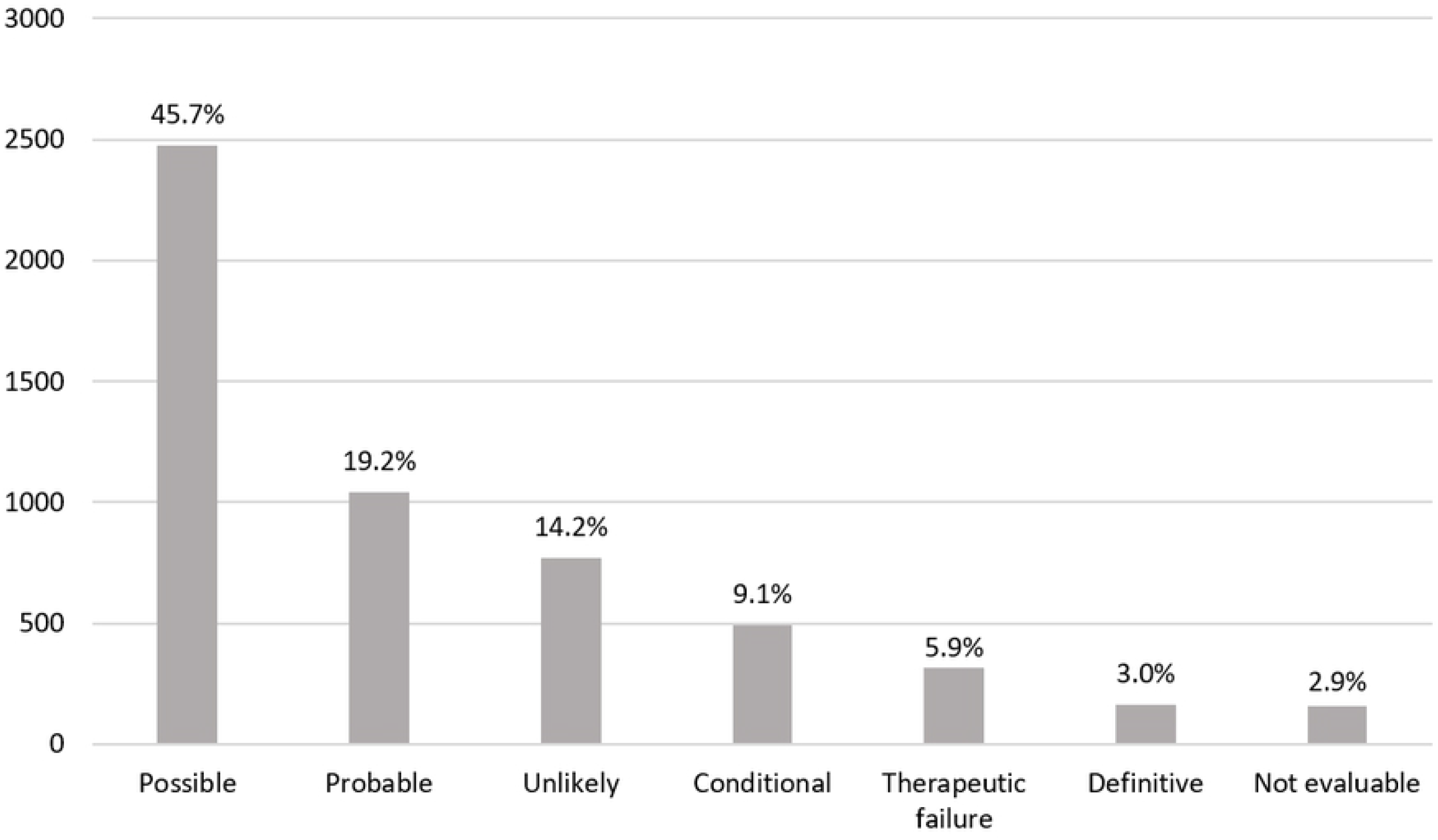
Adverse drug reactions by biotech agents according to probability classification to the World Health Organization in patients of Colombia.

**Table 4.**
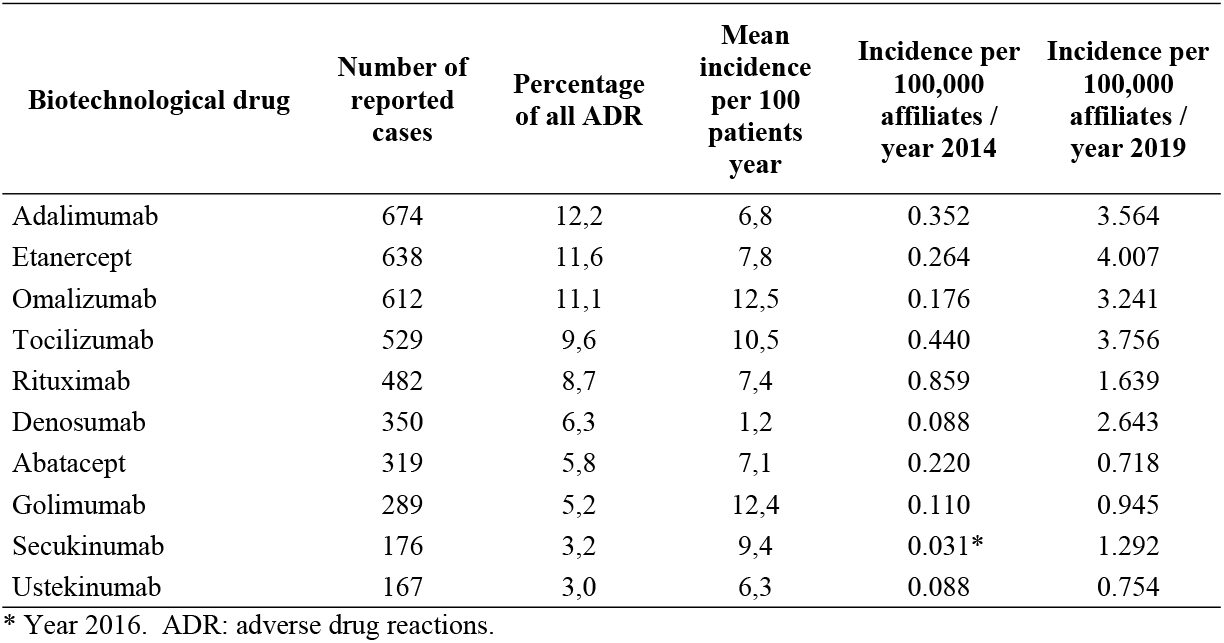
Top 10 of the biological agents with the highest number of reports and their incidences in Colombia 2014-2019

| Biotechnological drug | Number of reported cases | Percentage of all ADR | Mean incidence per 100 patients year | Incidence per 100,000 affiliates / year 2014 | Incidence per 100,000 affiliates / year 2019 |
| --- | --- | --- | --- | --- | --- |
| Adalimumab | 674 | 12,2 | 6,8 | 0.352 | 3.564 |
| Etanercept | 638 | 11,6 | 7,8 | 0.264 | 4.007 |
| Omalizumab | 612 | 11,1 | 12,5 | 0.176 | 3.241 |
| Tocilizumab | 529 | 9,6 | 10,5 | 0.440 | 3.756 |
| Rituximab | 482 | 8,7 | 7,4 | 0.859 | 1.639 |
| Denosumab | 350 | 6,3 | 1,2 | 0.088 | 2.643 |
| Abatacept | 319 | 5,8 | 7,1 | 0.220 | 0.718 |
| Golimumab | 289 | 5,2 | 12,4 | 0.110 | 0.945 |
| Secukinumab | 176 | 3,2 | 9,4 | 0.031* | 1.292 |
| Ustekinumab | 167 | 3,0 | 6,3 | 0.088 | 0.754 |

\* Year 2016. ADR: adverse drug reactions.

## Discussion

It was possible to determine which drugs of biotechnological origin were most frequently involved in ADRs in the Colombian population; this is the objective of this study. Biotech drugs must have specific pharmacovigilance considerations, including closer monitoring that can ensure their effectiveness and safety [9]. Although biotech drugs are less commonly used than synthetic drugs (approximately 20% of drugs currently are biotech drugs), they are very often associated with adverse events, some of which are serious and even lethal [10, 11]. In recent years, reports of biotech drugs associated with ADRs have increased worldwide, as also observed in this study, which is perhaps related to greater notification by prescribing physicians, nurses and patients and the greater use of these drugs for the treatment of a large number of pathological entities [4, 12].

ADRs associated with biotech drugs occur more frequently in women, as documented in other studies conducted in Spain (82.9%) [13], the United States (75.5%) [14] and Italy (54.3-71.3%) [4, 15], in agreement with the present finding. This is probably because many of the pathologies for which biotech drugs are used have a known predominance in women, including autoimmune diseases such as rheumatoid arthritis [16, 17] or oncological diseases [18], which expose women to a greater probability of use and of developing ADRs. In the present study, antineoplastics and immunomodulators were the biotech drugs most frequently associated with this type of event, in agreement with the findings of Cutroneo et al. in Italy [4].

The ADRs most documented in different studies are those related to infections [14, 15, 19–21], general manifestations or those associated with the administration of biotech drugs [12, 22, 23] and those related to the skin or subcutaneous tissues [4, 12, 23, 24]. However, the present study found that the most common ADRs were those related to the respiratory tract, diverging with what was found in other studies where their frequency was much lower (16.8 vs. 3.8-10.8%) [4, 12, 24], probably because we included in this category infections such as pneumonia and bronchitis, among others, which increased the proportion of respiratory tract-related ADRs.

Type A and B ADRs were the most frequent in a previous study conducted in Colombia [25]. That study analyzed a cohort of patients with rheumatoid arthritis treated with synthetic (sDMARD) and biologic (bDMARD) disease-modifying antirheumatic drugs and found that of all ADRs, 87.7% were type A (sDMARD: 70.2% vs. bDMARD: 17.5%) and 12.3% were type B (sDMARD: 8.1% vs. bDMARD: 4.2%); no other ADR types were observed [25]. In addition, according to severity, 22.6% of the reports were classified as serious, consistent with what was found in Italy (9.8-25.5%) [15, 23], Japan (18.5-23.4%) [20, 26], Spain (21.7%) [19], Brazil (25.0%) [22] and Korea (32.3%) [12]. Among severe reactions, the possibility of developing cancer, infections, hypersensitivity reactions and major cardiovascular events is described in the literature [10, 12, 14, 19, 22], and fatalities can also occur, which in this report corresponded to 0.1% of all ADRs, a rate lower than that documented in another study [27]. Death does not correspond to an adverse event but rather to a fatal outcome that can also be explained by the underlying disease of the patient. The classification of the severity of the event is independent of the degree of association with its causality.

In the present study, the main biotech drugs related to ADRs were adalimumab and etanercept, in agreement with other studies [13, 15, 22], but the incidence per 100 patients per year was higher than that reported in Spain in patients with rheumatoid arthritis (8.1 for adalimumab and 5.1 for etanercept) [19]. The incidence in our study was determined in a general manner for all ADRs, while the Spanish study only considered serious ADRs [19]. In Brazil, in patients with rheumatoid arthritis and psoriatic arthritis, 55.2% and 19.8% of ADRs were secondary to adalimumab and etanercept, respectively [22]. In Spain, in patients with rheumatoid arthritis, 35.1% and 21.6% of ADRs were due to adalimumab and etanercept, respectively [13]. In Italy, Barbieri et al. studied patients with inflammatory arthritis and found that 27.3% and 19.0% of ADRs were due to etanercept and adalimumab, respectively [15]. However, many studies also report a high proportion of ADRs secondary to infliximab [3, 19, 27], which was not observed in the present study due to the low use of this drug in Colombia. Additionally, the proportion of ADRs secondary to omalizumab in the present study is noteworthy. In Kuwait, in patients with asthma treated with omalizumab, 34.3% had ADRs, and 42.8% discontinued treatment [28].

One of the limitations of this study is its observational nature, as it is based on a database of reports that does not include variables such as patient age, comorbidities or comedications. Additionally, the proportion of patients who had to discontinue treatment due to an ADR was not analyzed, nor were the time elapsed from the administration of the biotech drug and to the onset of the ADR, concomitant medications or ADRs associated with previous treatments, although all of these factors are identified in the individual report and monitoring of each case. Moreover, for this analysis, no distinction was made between innovative and biosimilar drugs. However, the strongest point of this study is that it compiled ADR reports from one of the largest cohorts of patients in Colombia, for which exhaustive follow-ups were performed to identify the causality association.

## Conclusions

Based on our findings, we conclude that the reporting of ADRs has increased in recent years and that the reactions are mostly classified as A or B, are categorized as serious in almost one-fifth of the reported cases and are mainly associated with immunomodulators and antineoplastic agents. It is important to empower physicians and entire health teams to improve the traceability of adverse reactions and thus optimize and strengthen pharmacovigilance programs. This type of study can support decision makers in aspects that benefit patient safety and interaction with health systems.

## Acknowledgments

Soffy Claritza López, for her work in obtaining the database.

## Declarations

### Declaration of interest

The authors declare no conflicts of interest.

### Funding sources

The present study did not receive funding.

### Availability of data and material

protocolos.io

### Code availability

dx.doi.org/10.17504/protocols.io.bkcfkstn

### Authors’ contributions

ALJM: conceptualization, methodology, validation, formal analysis. YCMY: conceptualization, methodology, validation, formal analysis. IYPF: data curation, review and editing. LFVR: investigation, data curation, writing original draft. JEMA: methodology, validation, formal analysis, resources, writing, review and editing, supervision.

### Author responsibility

The corresponding author confirm full access to all data in the study and final responsibility.

## References

1. Figueredo JLS, Bautista SC, Barrenechea LM, Ronsano JBM. Eficiencia de los fármacos de origen biotecnológico en el marco terapéutico actual, según los estudios farmacoeconómicos disponibles. PharmacoEconomics Spanish Research Articles. 2008;5(4):119–33.

2. Machado-Alba JE, Moncada-Escobar JC. Reacciones adversas medicamentosas en pacientes que consultaron a instituciones prestadoras de servicios en Pereira, Colombia [Adverse drug reactions in patients attending in emergency service]. Rev Salud Publica (Bogota). 2006;8(2):200–208.

3. Vermeer NS, Giezen TJ, Zastavnik S, Wolff-Holz E, Hidalgo-Simon A. Identifiability of Biologicals in Adverse Drug Reaction Reports Received From European Clinical Practice. Clin Pharmacol Ther. 2019;105(4):962–969.

4. Cutroneo PM, Isgrò V, Russo A, et al. Safety profile of biological medicines as compared with non-biologicals: an analysis of the italian spontaneous reporting system database. Drug Saf. 2014;37(11):961–970.

5. Aspden P, Corrigan J, Wolcott J, Erickson S. Patient safety: Achieving a New Standard for Care. Washington. National Acedemies Press. 2004. Pp. 200–24

6. Heather EM, Payne K, Harrison M, Symmons DP. Including adverse drug events in economic evaluations of anti-tumour necrosis factor-α drugs for adult rheumatoid arthritis: a systematic review of economic decision analytic models. Pharmacoeconomics. 2014;32(2):109–134.

7. Sultana J, Cutroneo P, Trifirò G. Clinical and economic burden of adverse drug reactions. J Pharmacol Pharmacother. 2013;4(Suppl 1):S73–S77.

8. Giezen TJ, Mantel-Teeuwisse AK, Leufkens HG. Pharmacovigilance of biopharmaceuticals: challenges remain. Drug Saf. 2009;32(10):811–817.

9. O’Callaghan J, Griffin BT, Morris JM, Bermingham M. Knowledge of Adverse Drug Reaction Reporting and the Pharmacovigilance of Biological Medicines: A Survey of Healthcare Professionals in Ireland. BioDrugs. 2018;32(3):267–280.

10. Sousa J, Taborda-Barata L, Monteiro C. Biological therapy-associated adverse reactions in asthma: analysis of reporting to the Portuguese pharmacovigilance system. Expert Opin Drug Saf. 2020;19(1):99–106.

11. Klein K, Scholl JH, Vermeer NS, Broekmans AW, Van Puijenbroek EP, De Bruin ML, et al. Traceability of Biologics in The Netherlands: An Analysis of Information-Recording Systems in Clinical Practice and Spontaneous ADR Reports. Drug Saf. 2016;39(2):185–192.

12. Sim DW, Park KH, Park HJ, Son YW, Lee SC, Park JW, et al. Clinical characteristics of adverse events associated with therapeutic monoclonal antibodies in Korea. Pharmacoepidemiol Drug Saf. 2016;25(11):1279–1286.

13. Abasolo L, Leon L, Rodriguez-Rodriguez L, Tobias A, Rosales Z, Maria Leal J, et al. Safety of disease-modifying antirheumatic drugs and biologic agents for rheumatoid arthritis patients in real-life conditions. emin Arthritis Rheum. 2015;44(5):506–513.

14. Ringold S, Hendrickson A, Abramson L, Beukelman T, Blier PR, Bohnsack J, et al. Novel method to collect medication adverse events in juvenile arthritis: results from the childhood arthritis and rheumatology research alliance enhanced drug safety surveillance project. Arthritis Care Res (Hoboken). 2015;67(4):529–537.

15. Barbieri MA, Cicala G, Cutroneo PM, Gerratana E, Palleria C, De Sarro C, et al. Safety Profile of Biologics Used in Rheumatology: An Italian Prospective Pharmacovigilance Study. J Clin Med. 2020;9(4):1227.

16. Ortona E, Pierdominici M, Maselli A, Veroni C, Aloisi F, Shoenfeld Y. Sex-based differences in autoimmune diseases. Ann Ist Super Sanita. 2016;52(2):205–212.

17. Ngo ST, Steyn FJ, McCombe PA. Gender differences in autoimmune disease. Front Neuroendocrinol. 2014;35(3):347–369.

18. Rojas K, Stuckey A. Breast Cancer Epidemiology and Risk Factors. Clin Obstet Gynecol. 2016;59(4):651–672.

19. Leon L, Gomez A, Vadillo C, Pato E, Rodriguez-Rodriguez L, Jover JA, et al. Severe adverse drug reactions to biological disease-modifying anti-rheumatic drugs in elderly patients with rheumatoid arthritis in clinical practice. Clin Exp Rheumatol. 2018;36(1):29–35.

20. Yamanaka H, Hirose T, Endo Y, Sugiyama N, Fukuma Y, Morishima Y, et al. Three-year safety and two-year effectiveness of etanercept in patients with rheumatoid arthritis in Japan: Results of long-term postmarketing surveillance. Mod Rheumatol. 2019;29(5):737–746.

21. Narongroeknawin P, Chevaisrakul P, Kasitanon N, Kitumnuaypong T, Mahakkanukrauh A, Siripaitoon B, et al. Drug survival and reasons for discontinuation of the first biological disease modifying antirheumatic drugs in Thai patients with rheumatoid arthritis: Analysis from the Thai Rheumatic Disease Prior Authorization registry. Int J Rheum Dis. 2018;21(1):170–178.

22. de Camargo MC, Barros BCA, Fulone I, Silva MT, Silveira MSdN, Camargo IAd, et al. Adverse Events in Patients With Rheumatoid Arthritis and Psoriatic Arthritis Receiving Long-Term Biological Agents in a Real-Life Setting. Front Pharmacol. 2019;10:965.

23. Roberti R, Iannone LF, Palleria C, De Sarro C, Spagnuolo R, Barbieri MA, et al. Safety profiles of biologic agents for inflammatory bowel diseases: a prospective pharmacovigilance study in Southern Italy. Curr Med Res Opin. 2020;1–7.

24. Palleria C, Iannone L, Leporini C, Citraro R, Manti A, Caminiti M, et al. Implementing a simple pharmacovigilance program to improve reporting of adverse events associated with biologic therapy in rheumatology: Preliminary results from the Calabria Biologics Pharmacovigilance Program (CBPP). PLoS One. 2018;13(10):e0205134.

25. Machado-Alba JE, Ruiz AF, Machado-Duque ME. Adverse drug reactions associated with the use of disease-modifying anti-rheumatic drugs in patients with rheumatoid arthritis. Rev Panam Salud Publica. 2014;36(6):396–401.

26. Koike T, Harigai M, Inokuma S, Inoue K, Ishiguro N, Ryu J, et al. Postmarketing surveillance of the safety and effectiveness of etanercept in Japan. J Rheumatol. 2009;36(5):898–906.

27. Ha D, Lee SE, Song I, Lim SJ, Shin JY. Comparison of signal detection of tumour necrosis factor-α inhibitors using the Korea Adverse Events Reporting System Database, 2005-2016. Clin Rheumatol. 2020;39(2):347–355.

28. Al-Ahmad M, Nurkic J, Maher A, Arifhodzic N, Jusufovic E. Tolerability of Omalizumab in Asthma as a Major Compliance Factor: 10-Year Follow Up. Open Open Access Maced J Med Sci. 2018;6(10):1839–1844.

